# Implementations of the chemical structural and compositional similarity metric in R and Python

**DOI:** 10.1101/546150

**Authors:** Asker Brejnrod, Madeleine Ernst, Piotr Dworzynski, Lasse Buur Rasmussen, Pieter C. Dorrestein, Justin J.J. van der Hooft, Manimozhiyan Arumugam

## Abstract

**Motivation:** Tandem mass spectrometry (MS/MS) has the potential to substantially improve metabolomics by acquiring spectra of fragmented ions. These fragmentation spectra can be represented as a molecular network, by measuring cosine distances between them, thus identifying signals from the same or similar molecules. Metrics that enable comparison between pairs of samples based on their metabolite profiles are in great need. Taking inspiration from the successful phylogeny-aware beta-diversity measures used in microbiome research, integrating chemical similarity information about the features in addition to their abundances could lead to better insights when comparing metabolite profiles. Chemical Structural and Compositional Similarity (CSCS) is a recently published similarity metric comparing the full set of signals and their chemical similarity between two samples. Efficient, scalable and easily accessible implementations of this algorithm is currently lacking. Here, we present an easily accessible and scalable implementation of CSCS in both python and R, including a version not weighted by intensity information.

**Results:** We provide a new implementation of the CSCS algorithm that is over 300 times faster than the published implementation in R, making the algorithm suitable for large-scale metabolomics applications. We also show that adding chemical information enriches existing methods. Furthermore, the R implementation includes functions for exporting molecular networks directly from the mass spectral molecular networking platform GNPS for ease of use for downstream applications.

**Availability:** github.com/askerdb/rCSCS, github.com/askerdb/pyCSCS

## 1 Introduction

Liquid chromatography tandem - mass spectrometry (LC-MS/MS) is gaining more and more popularity in the metabolomics field with a wide range of applications (e.g. [11, 9, 13, 12]). As an extension of LC-MS, one of the most frequently used analytical platforms in mass spectrometry-based metabolomics [27, 4], LC-MS/MS does not only provide a high coverage and sensitivity towards semi-polar metabolites, but also provides chemical structural information of the metabolites investigated. In LC-MS/MS, metabolites are ionized and fragmented and the resulting fragmentation fingerprints of mass-to-charge ratios (m/z) are characteristic to the molecular structure of the metabolite. Furthermore, metabolites resulting in similar fragmentation patterns presumably exhibit similar chemical structures [24, 25]. The similarity of these fingerprints can be calculated using the cosine score and represented as a graph, resulting in so-called mass spectral molecular networks (recently reviewed in [14]). This approach, popularized by the user-friendly Global Natural Products Social Molecular Networking (GNPS) platform [24], has been highly effective in visualizing structural chemical relatedness across samples as well as in aiding the structural identification of metabolites. Several studies have focused on deriving and applying new distance metrics to metabolomics profiles to compare pairs of samples [16, 15, 8]. Among these, the chemical structural and compositional similarity (CSCS) as proposed by Sedio and collaborators [19, 17, 18], accounts for the chemical structural similarity across metabolites by integrating the similarity of their MS/MS fragmentation patterns through the cosine score. Even though metabolites can be identified for only about 2 % of all signals in a typical LC-MS/MS experiment [1], CSCS has the advantage to integrate chemical structural information without depending on metabolite identification, thus enabling a comparison of the entire range of detected molecules. CSCS can be used to visualize chemical differences across samples by using Principal Coordinate Analysis (PCoA) plots [5] or distance based hypothesis testing of differences. Here, we implement CSCS in user-friendly python and R packages, and demonstrate its performance by computing commonly used metrics on nearly 500 LC-MS/MS samples of the American Gut Project [9] as well as fecal samples obtained from children with Crohn’s disease at different time points during nutritional therapy [22].

## 2 Methods

### 2.1 Mathematical exposition

Consider two samples A and B with the union of features [1, …, *C*], and intensity vectors *A* = [*I*_*a*1_, …, *I*_*aC*_] and *B* = [*I*_*b*1_, …, *I*_*bC*_]. Calculation of the Chemical Structural and Compositional Similarity (CSCS) takes the following inputs: a chemical similarity matrix, *CSS*, and the two vectors of ion intensities, *A* and *B*. *CSS* is defined as a matrix of pairwise cosine distances

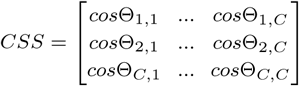

In practice this matrix can be calculated from a file specifying pairwise cosine distances across nodes in a mass spectral molecular network, which can be downloaded from GNPS.

The CSCS as defined by Sedio and collaborators [19], referred to here as weighted CSCS (*CSCSw*) is then defined as:

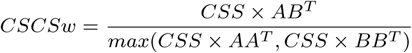

where × is element-wise multiplication.

Analogously, we here define the unweighted *CSCSu* where all elements of *A* and *B* are dichotomized into 1 or 0 upon presence or absence of a signal in the sample, respectively. CSCS is a metric of similarity. However, many downstream analyses within metabolomics rely on a dissimilarity matrix (e.g. PCoA), thus our libraries return a dissimilarity matrix, corresponding to 1-CSCS.

### 2.2 Implementation

Implementations are available at https://github.com/askerdb/ rCSCS as an R [21] package, which can be installed through devtools [26], and at https://anaconda.org/askerdb/pycscs as a python package, which can be installed through conda. The R package depends on the packages foreach [2], igraph [3] and Rcurl[7], wheras the python package depends on numpy [23], pandas [10], scikit bio (http://scikit-bio.org) as well as the sparse matrix implementation in scipy [6].

### 2.3 Data

Run time benchmarks were computed on 18 fecal samples obtained from children with Crohn’s disease at different time points during nutritional therapy [22]. This dataset is publicly available at https://massive.ucsd.edu/ under the MassIVE accession number MSV000081120. A total of 1245 MS/MS features were retrieved for this dataset using the GNPS networking parameters publicly accessible at https://gnps.ucsd.edu/ProteoSAFe/status. jsp?task=b0524246804a4b50a8a4ec6244a8be2e. Samples from the American Gut Project data are publicly available under the MassIVE accession number MSV000080179 [9]. This dataset consisted of 489 samples and 16349 features using GNPS networking parameters publicly accessible at https://gnps.ucsd.edu/ProteoSAFe/ status.jsp?task=a07557dc26cc4d3f8a2076d5ae0898a2.

## 3 Results

### 3.1 Benchmarks

We computed CSCS for a dataset consisting of 18 fecal samples obtained from children with Crohn’s disease at different time points during nutritional therapy [22] and a total of 1245 MS/MS features, a realistically sized dataset on a single core of a Macbook Pro (2.8 GHz i7), and compared it against the published implementation from Sedio and collaborators [19]. Results shown in Table 1 include run time overhead that should be minimal in realistic applications. To demonstrate the utility of this implementation on a scale that is relevant to high-throughput collection of data, we used the American Gut Project that consists of 489 samples with 16349 MS/MS features when downloaded from GNPS. Analysis of this dataset with our python implementation finished in 7.5 hours on 40 CPU cores. We did not compare run time of the original implementation [19], as this will likely not finish in a reasonable time.

**Table 1.**
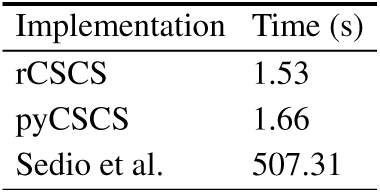
CSCS timing benchmarks for a dataset consisting of 18 fecal samples obtained from children with Crohn’s disease at different time points during nutritional therapy [22].

### 3.2 Chemical information produces distinct similarities

To evaluate how the CSCS metrics compare with other popular metrics, we compared them with Bray-Curtis, Jaccard and Canberra metrics estimated for a dataset of 594 metabolome samples from the American Gut Project [9]. We used Procrustes analysis[20] to measure the similarity between the distance matrices and used this information as input for a PCoA plot (Figure 1). Metrics with chemical information are clearly separated from those without, and on this dataset the weighting of chemical information with ion intensities drives the separation on the axis that explains the most variance, while the unweighted CSCS is distinctly separated on the second principal component.

**Fig. 1.**
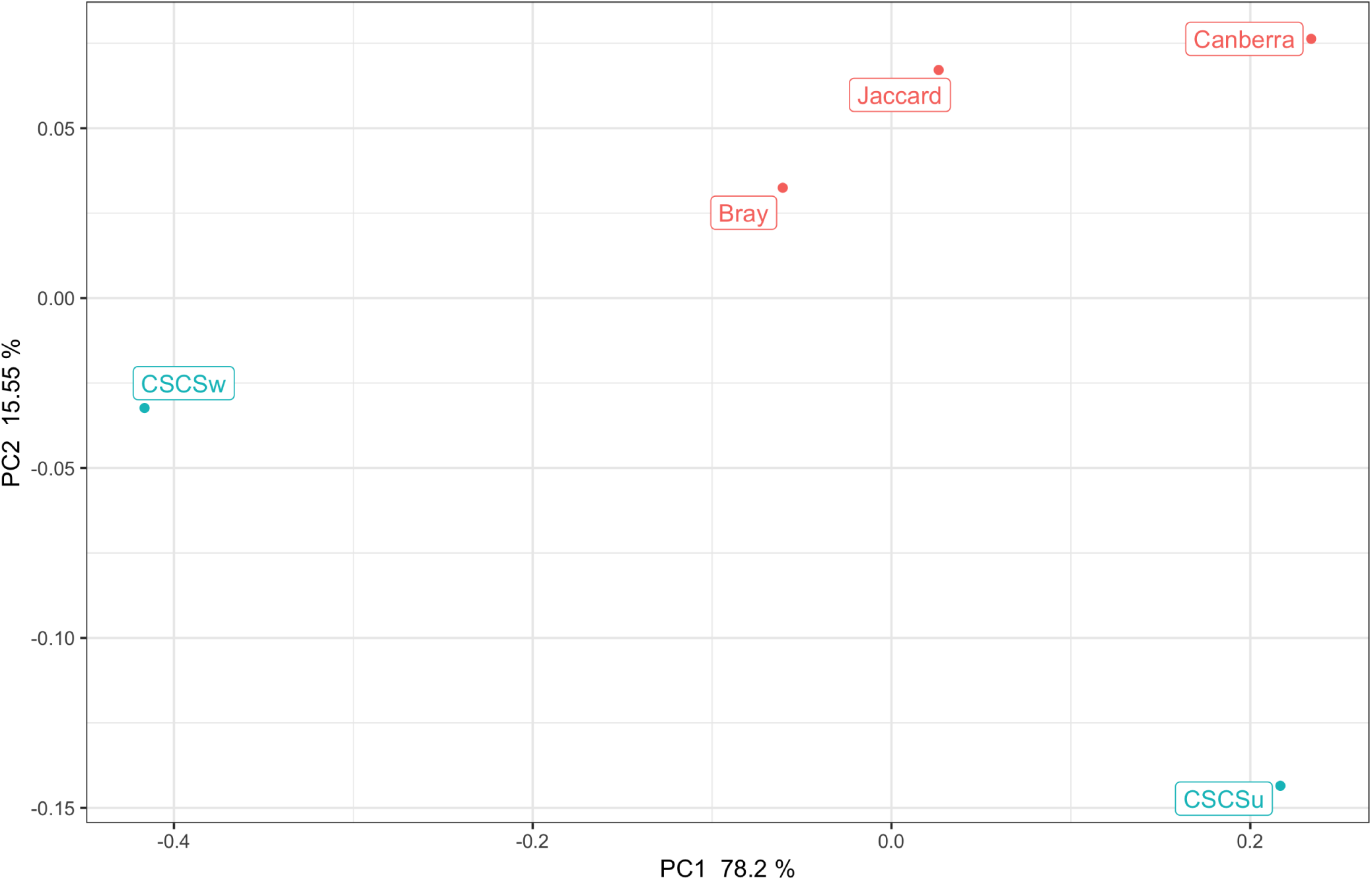
PCoA of the procrustes distances between various distance metrics commonly used in metabolomics. Metrics colored in green include chemical information, either weighted by ion intensities or unweighted. Metrics with no chemical information are colored red. information

## 4 Conclusion

We have implemented the calculation of weighted and unweighted CSCS distances in R and Python. Through benchmarking of publicly available data we have demonstrated the highly significant run time improvement, which expands the application of CSCS to large datasets.

## Acknowledgements

We would like to acknowledge Brian Sedio and collaborators for proposing and implementing the original chemical structural and compositional similarity metric, and making software available for direct comparisons.

